# Inhibition of EMT driver PTK6 enhances anti-tumor immune responses against triple-negative breast cancer

**DOI:** 10.1101/2025.01.11.632044

**Authors:** Ibuki Harada, Criseyda Martinez, Koichi Ito, Eunjee Lee, Jun Zhu, Hanna Y Irie

**Affiliations:** Division of Hematology and Medical Oncology, Department of Medicine, Icahn School of Medicine at Mount Sinai, New York, NY 10029, USA; Sema4, Stamford, Connecticut; Department of Oncological Sciences, Icahn School of Medicine at Mount Sinai, New York, New York, NY 10029, USA

## Abstract

The non-receptor tyrosine kinase PTK6 is expressed in 70% of triple negative breast cancers (TNBC) and is an oncogenic driver of epithelial-mesenchymal transition (EMT). EMT promotes metastasis and immune evasion of TNBC. Therefore, targeting EMT drivers could reverse these properties and lead to more favorable outcomes. Treatment of TNBC tumors with a small molecule inhibitor of PTK6 kinase (P21d) suppressed their growth *in vivo*. Tumor inhibition by P21d is dependent on an induced immune response because: 1) inhibition is observed in immunocompetent, but not immunodeficient, mice; 2) P21d increases tumor-infiltrating CD8^+^ T and NK cells and decreases immunosuppressive myeloid-derived suppressor cells; and 3) tumor inhibition by P21d is abrogated by co-treatment with NK or CD8^+^ T cell-depleting antibodies. These effects on tumor growth and cytotoxic TILs are phenocopied by the knockdown of tumoral PTK6 or SNAIL, which supports EMT inhibition as a mechanism for enhanced anti-tumor immune response. RNA sequencing (RNA-seq) profiling of P21d-treated tumors also revealed changes consistent with activation of the immune response and identified CXCL10 as a critical chemokine induced intratumorally by P21d that promotes recruitment of NK/CD8+ T cells to the tumor site, leading to tumor growth inhibition. Our study highlights the novel tumor immune microenvironmental functions of PTK6 with important consequences for tumor growth that could lead to new immunotherapeutic approaches for TNBC.

## INTRODUCTION

Triple-negative breast cancer (TNBC), which constitutes approximately 15% of all diagnosed breast cancers, remains the most aggressive breast cancer subtype and is associated with a higher probability of metastasis and early recurrence compared to other breast cancer subtypes [1, 2]. TNBC is characterized by the lack of expression of estrogen receptor (ER), progesterone receptor (PR), and human epidermal growth factor receptor 2 (HER2) [3]. Despite multimodality treatment consisting of systemic therapies, surgery and radiation, many patients still experience early relapses with metastatic disease. Therefore, novel therapeutic targets and strategies to enhance the efficacy of current treatments may help alleviate the significant morbidity and mortality associated with recurrence and metastatic disease.

Epithelial-mesenchymal transition (EMT), a process by which epithelial cells assume mesenchymal markers and phenotypes such as increased migration and invasion, has been closely linked to chemotherapy resistance and metastasis [4–9]. In addition, EMT promotes immune evasion and immunosuppressive tumor microenvironment [9]. For example, mesenchymal-like tumor cells inhibit the function and recruitment of NK and CD8^+^ T cells by upregulating immune checkpoint molecules such as PD-L1 and producing soluble factors such as CCL2, CXCL1/2 and CSF1, which activate the function and recruitment of suppressive cells known as Myeloid derived suppressor cells (MDSCs), regulatory T cells (Tregs) and tumor-associated macrophages (TAM) [10–15]. Hence, targeting EMT could enhance sensitivity to chemotherapy and immunotherapy for more complete eradication of tumor cells and prevention of metastases.

We previously identified Protein tyrosine kinase 6 (PTK6) as an EMT driver that is highly expressed in approximately 40% of TNBCs [16]. PTK6 is a non-receptor tyrosine kinase that is expressed in several cancers, including breast ovarian, and prostate cancers, with low to absent expression in the normal tissue counterparts [17, 18]. PTK6 expression is associated with prognostic significance in breast cancer and promotes growth and survival of cancer cells. PTK6 drives EMT and promotes the invasion and metastases of TNBC by stabilizing the expression of SNAIL, a transcriptional co-repressor of E-cadherin[16]. PTK6 regulates SNAIL ubiquitination and proteasome-dependent degradation by modulating its interaction with MARCH2 E3 ligase in a PTK6 kinase activity-dependent manner [19].

The interaction between cancer cells that have undergone EMT, and their tumor immune microenvironment may also determine their sensitivity to various immunotherapies [20–23]. Immune cell infiltration and the composition of immune infiltrates are particularly critical for triple negative breast cancer, with both prognostic and predictive significance [24–26]. Increased cytotoxic TILs observed in patient TNBCs are associated with better outcomes and response to chemo/immunotherapy.

Analysis of the Cancer Genome Atlas (TCGA) revealed that PTK6 is expressed in many malignancies, and that PTK6 expression has prognostic significance with higher levels associated with worse outcomes [27]. PTK6 expression was reported to be inversely correlated with distinct immune signatures, including those of cytotoxic immune cells [27]. PTK6 has also been proposed as a biomarker of immunotherapy response in renal cell carcinoma [28]. Given PTK6’s role as an EMT driver and as a TNBC oncogene, we reasoned that PTK6 could play a critical role in regulating the immune microenvironment of TNBC and regulating anti-tumor immune responses that can impact outcomes and/or therapeutic response.

In this study, we assessed the functions of PTK6 in regulating the tumor immune microenvironment of TNBC, the underlying mechanisms of immunomodulatory effects and the impact of PTK6-dependent immune regulation on TNBC tumor growth control.

## MATERIALS AND METHODS

### Antibodies and Reagents

Antibodies for flow cytometry were purchased from Biolegend: CD45 (Clone:30-F11, RRID:AB_10899570 and RRID:AB_312980), F4/80 (Clone:BM8, RRID:AB_893478 and RRID:AB_893484), CD3e (Clone:500A2, RRID:AB_2632668), CD49b (Clone:HMa2, RRID:AB_313029), CD11c (Clone:N418, RRID:AB_830646), NK1.1 (Clone:S17016D, RRID:AB_3083138), NKp46 (Clone:29A1.4, RRID:AB_2298210), CD11b (Clone:M1/70, RRID:AB_312788 and RRID:AB_312790), Ly6G/Ly6C (Clone:RB6-8C5, RRID:AB_2562214), CD8a (Clone:53-6.7, RRID:AB_312747), CD4 (Clone:GK1.5, RRID:AB_312690), Granzyme B (Clone:QA16A02, RRID:AB_2687029) and Isotype control IgG (BioLegend, RRID:AB_11203529). The following antibodies were used for Western blot analysis: phospho-PTK6 (Millipore Sigma, RRID: AB_612089), GAPDH (Clone, Cell Signaling cat# or RRID number), SNAIL (Cell Signaling Technology, RRID:AB_2255011), PTK6 (Sigma-Aldrich, RRID:AB_10639478)), and PTK6 (Santa Cruz Biotechnology, RRID:AB_2174225).

CD8 and NK cell depleting antibodies were purchased from Biolegend (RRID:AB_2810323 and RRID:AB_2800567). Dimethyl sulfoxide (DMSO) was purchased from Sigma-Aldrich. [4-[[6-cyclopropyl-3-(1H-pyrazol-4-yl)imidazo[1,2-a]pyrazin-8-yl]amino]-3-fluorophenyl]-morpholin-4-ylmethanone (P21d) was purchased from Tocris.

For immunofluorescence, the CD8 antibody was purchased from Abcam (RRID: AB_2860566). The NKp46 antibody was purchased from BioLegend (RRID:AB_2298210). Secondary antibody was purchased from Invitrogen (RRID:RRID:AB_2535850).

### Cell lines and culture

MMTV-myc cells were obtained from Dr. Eduardo Farias (Icahn School of Medicine at Mount Sinai, New York, NY), and cultured in DMEM/F12 supplemented with 5% FBS and 4 mg/mL insulin as previously described [16]. 4T-1 cells were obtained from ATCC and cultured according to ATCC guidelines (RRID:CVCL_0125). PTK6-overexpressing MCF10A cells were generated by viral infection and were cultured as previously described [16].

### Mice and tumor cell transplantation

Six-week-old female FVB/N mice (Charles River Laboratories, RRID:IMSR_JAX:036633), NOD.Cg-PrkdcscidIl2rgtm1Wjl/SzJ (NSG) mice (Jackson Laboratories, RRID:IMSR_JAX:005557) and BALB/C mice (Charles River Laboratories, RRID:IMSR_JAX:000651) were purchased and maintained in a pathogen-free facility. Tumor cells (1 × 10^5^) were injected into the right inguinal mammary fat pad of mice. P21d was administered via intraperitoneal injection. All animal procedures were approved by the Institutional Animal Care and Use Committees (IACUCs) of the Icahn School of Medicine at Mount Sinai.

### Viral transduction and tumor cell transplantation

Viral particles containing retroviral or lentiviral vectors encoding cDNA or shRNA sequences were generated by co-transfection of 293T cells with vector and packaging plasmids (Delta 8.9 and pCMV-VSV-G, RRID:Addgene_8454) using Lipofectamine 2000 and Plus Reagent (Life Technologies). Viral supernatants were collected and used to infect cells.

### RNA-sequencing analysis

Tumor RNA samples from 7 days P21d-treated MMTV-myc transplanted FVB/NJ mice (n = 3) and these controls (n = 3) underwent RNA sequencing (RNA-seq) at the New York Genome Center. The New York Genome Center adapted a previously reported method [29, 30]. Briefly, the RNA-seq libraries were used for the tag-mentation reaction carried out using the Illumina TruSeq Stranded mRNA Sample Preparation Kit (Illumina) according to the manufacturer’s instructions. Agilent Bioanalyzer and quantification of the final library was performed using PicoGreen (fluorescence) methods. A single peak between 250 and 350 bp indicated a properly constructed and amplified library that was ready for sequencing. Sequencing was performed on a HiSeq 2500 using v4 SBS chemistry according to the Illumina protocol, as previously described [29, 30]. All samples were confirmed to be suitable for sequencing based on the size of the indexed cDNA library. FASTQ-formatted RNA-seq data were mapped onto the mouse reference genome GRCmm10 by using the STAR software (RRID:SCR_004463). The TPMs were calculated using STAR. For gene ontology (GO) analysis, differentially expressed genes (DEGs) were identified using the edgeR package (R version 3.1.2. RRID:SCR_012802) with FDR < 0.01 and logFC > 2 in samples between tumors treated with vehicle control vs. P21d. The list of DEGs was applied to the BiNGO App of Cytoscape (RRID:SCR_003032) as a query. GO terms enriched in this DEG list were ranked by *p*-value, and the top five enriched terms were listed.

Differentially expressed genes and enrichment of immune genes were identified using DESeq2 (PMID: 25516281). Genes with FDR ≤ 0.05 were deemed differentially expressed, with fold change <1 implying under-expression and vice versa. We identified 807 genes that were up-regulated and 1237 genes that were down-regulated with P21d treatment compared with the control. To identify the immune cells associated with these differentially expressed genes, we assessed the enrichment of immune marker genes by using Fisher’s exact test. Immune marker genes were defined based on the reference gene signature matrix LM22 (PMID: 25822800). LM22 is a signature matrix file consisting of 547 genes that distinguishes 22 mature human hematopoietic populations isolated from peripheral blood or *in vitro* culture conditions, including seven T cell types, naïve and memory B cells, plasma cells, NK cells, and myeloid subsets (PMID: 25822800). Immune cell markers were selected as the set of genes that had the highest signature values for each cell type. The expression levels of immunerelated genes were converted to Z-scores and are shown with a heatmap.

### Flow cytometric analysis

Cells were stained with 7-AAD (7-amino-actinomycin D) Viability Staining Solution (Biolegend, cat #420404) to detect and exclude dead cells. Intracellular staining was performed as previously described [31]. Briefly, TILs were fixed with 4% paraformaldehyde, treated with permeabilizing solution (50 mM NaCl, 5 mM EDTA, and 0.5% Triton X-100), and incubated with antibodies. Flow cytometry was performed using FACS Canto II and FACS Fortessa (BD Biosciences, RRID:SCR_018056 and RRID:SCR_018655), and data were analyzed using FlowJo software (FlowJo, LLC, RRID:SCR_008520).

### Western Blotting and Protein array

To prepare protein samples for Western Blotting or protein arrays, tumor cells were lysed with radioimmunoprecipitation assay (RIPA) buffer (Cell Signaling, Denver, MA). The cell lysates were centrifuged at 13,700 × g for 20 min at 4°C, and the supernatant was collected for subsequent western blot analysis. The concentrations of cell lysates were measured using the bicinchoninic acid (BCA) assay. The cell lysates were loaded by SDS-PAGE, transferred an Immobilon PVDF or nitrocellulose membrane and incubated with primary antibody overnight at 4 °C. After secondary antibody incubation, the membranes were developed using ECL Western Blotting Substrate (ThermoFisher Scientific or MiliporeSigma).

For the protein array, according to manufacturer’s instructions, cytokines and chemokines in the supernatant from MMTV-myc cell cultures were detected using Proteome Profiler array kits (R&D Systems, mouse: cat. #ARY028). Supernatants were collected from MMTV-myc cells after 48 hr of P21d treatment or shRNA-transduced cells.

### Migration assay of NK cells and CD8^+^ T cells

The migratory capacity of NK cells and CD8^+^ T cells was measured in 24-well modified chambers with 6.5 μm pore size polycarbonate membranes (Corning Inc., Corning, NY, USA). NK cells or CD8^+^ T cells isolated from mouse spleen were seeded in the upper chamber of the inserts at a density of 4 × 10^4^ cells per well in 300 μl serum-free medium. 1 mL of conditioned medium (culture supernatant) from MMTV-myc cells treated in vitro with P21d or vehicle control was added to the lower chamber. In the neutralization assay, an anti-CXCR3 (BioLegend, RRID: AB_11147756) or IgG1 isotype control (BioLegend, RRID:AB_11203529) antibody was added to the cells in the upper chamber at a concentration of 1 μg/ml 30 min before the addition of conditioned medium to the lower chamber. After 4 hours, migrated cells were visualized and counted.

### Immunofluorescence

Tumors recovered from control or P21d-treated FVB/N mice were frozen in O.C.T compound (ThermoFisher). Frozen sections (5 μm) were prepared and stored at -80°C. Sections were fixed with 4% para-formaldehyde/PBS for 20 minutes, followed by permeabilization with ice-cold 95% methanol for 30 minutes. Sections were washed with PBS, blocked with 3% BSA/PBS/0.1% TritonX for 1 hour at room temperature, and incubated with anti-CD8 and/or anti-NKp46 (1:100 dilution). After incubation at 4°C, sections were washed, incubated with AlexaFluor 555-conjugated secondary antibody (1:100 dilution) for 1 hour at room temperature, and mounted with VECTASHELD anti-fade media with DAPI (VectorLabs). Images were captured using a SP5 confocal microscope (Leica Microsystems). The gain and offset setting were fixed across all samples.

### Statistical analysis

Statistical significance of data was analyzed using the unpaired t test or ANOVA, after testing for normal distribution, unless indicated otherwise. P value is shown as: *, P < 0.05; **, P < 0.01; and ***, P < 0.001.

## RESULTS

### PTK6 inhibition suppresses TNBC tumor growth in immunocompetent mice

We determined the effect of inhibiting PTK6 on primary TNBC tumor growth in both immunocompetent and immunodeficient backgrounds using P21d, a small molecule inhibitor of PTK6 kinase activity that we previously validated [16]. P21d inhibits PTK6-dependent autophosphorylation, as well as phosphorylation of its targets and causes degradation of SNAIL, a co-repressor of E-cadherin and transcriptional driver of epithelial-mesenchymal transition (EMT) [16, 19] (**Fig. 1 A, D and Supplemental Fig S1**). Treatment of immunocompetent FVB/N mice bearing triple negative MMTV-myc tumors with P21d significantly inhibited tumor growth (**Fig. 1 B, C**). In contrast, P21d treatment did not inhibit the growth of the same MMTV-myc tumors implanted in immunodeficient NOD/SCID/IL-2Rψ− deficient mice (NSG) mice (**Fig. 1E, F**). These contrasting results are not due to differential amounts of intratumoral drug accumulation as evidenced by: 1) comparable suppression of SNAIL, a downstream target of PTK6 (**Fig. 1A, D**) and 2) mass spectrometry analysis of P21d-treated MMTV-myc tumors recovered from FVB/N or NSG mice showing higher levels of intratumoral P21d in NSG tumors (**Supplemental Fig S2**). These results suggest that tumor immune microenvironmental effects may mechanistically contribute to the tumor inhibitory effects of PTK6 inhibition.

**Figure 1.**
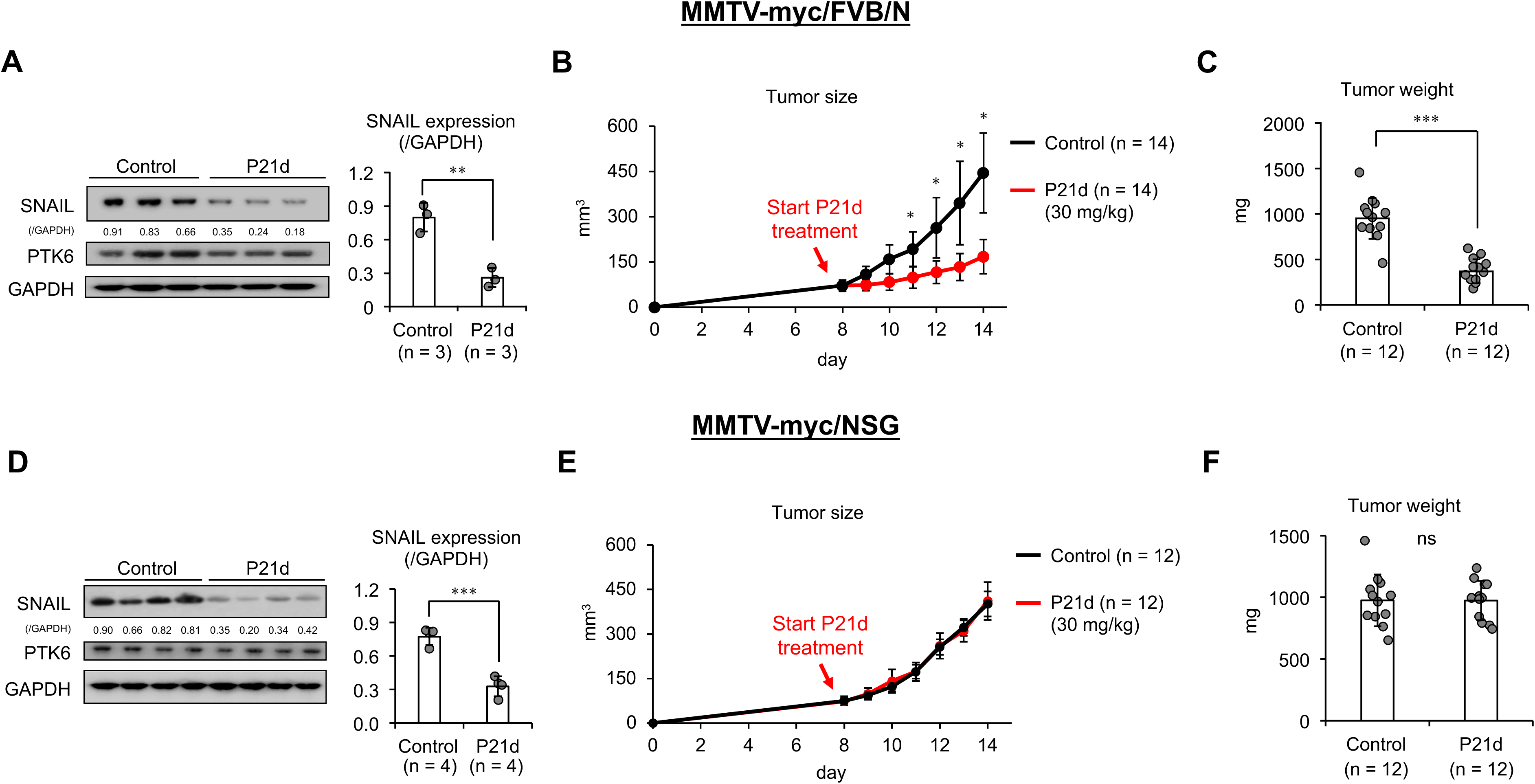
Effect of PTK6 kinase inhibitor treatment against murine TNBC. (A) MMTV-myc tumor bearing FVB/N mice were treated with vehicle control or P21d (30mg/kg/d) for three days. Tumors were recovered and SNAIL expression was assessed by Western blot. (n=3 mice/treatment group). (B,C) MMTV-myc tumor bearing FVB/N mice were treated with vehicle or P21d (30mg/kg) once primary tumors reached 100mm^3^ (n=14 mice/treatment group). Endpoint tumor weights were measured at day 14. (D) MMTV-myc tumor bearing NSG mice were treated with vehicle control or P21d (30mg/kg/d) for three days. Tumors were recovered and SNAIL expression was assessed by Western blot. (n=4 mice/treatment group). (E, F) MMTV-myc tumor bearing NSG mice were treated with vehicle or P21d (30mg/kg) once primary tumors reached 100mm^3^ (n=12 mice/treatment group). Endpoint tumor weights were measured at day 14. Data are the mean ± SD. \*\**p*=0.01, \*\*\**p*=0.0001 by Student’s *t*-test.

### PTK6 inhibition increases tumor-infiltrating cytotoxic immune cells

The differing effects of P21d treatment on TNBC tumors implanted in immunocompetent vs. immunodeficient mice suggest that immune regulation is critical for P21d-dependent tumor growth inhibition. This hypothesis was also supported by whole transcriptome sequencing (RNA-seq) analysis of MMTV-myc tumors extracted from FVB/N mice treated with vehicle control or P21d (**Fig. 2A**). Gene Ontology (GO) analysis of differentially expressed genes (DEGs) revealed an enrichment for genes related to positive regulation and activation of immune responses in P21d-treated tumors (**Fig. 2B**). Genes that were downregulated by P21d treatment relative to control were enriched in development and cell proliferation-related genes (**Fig. 2B**).

**Figure 2.**
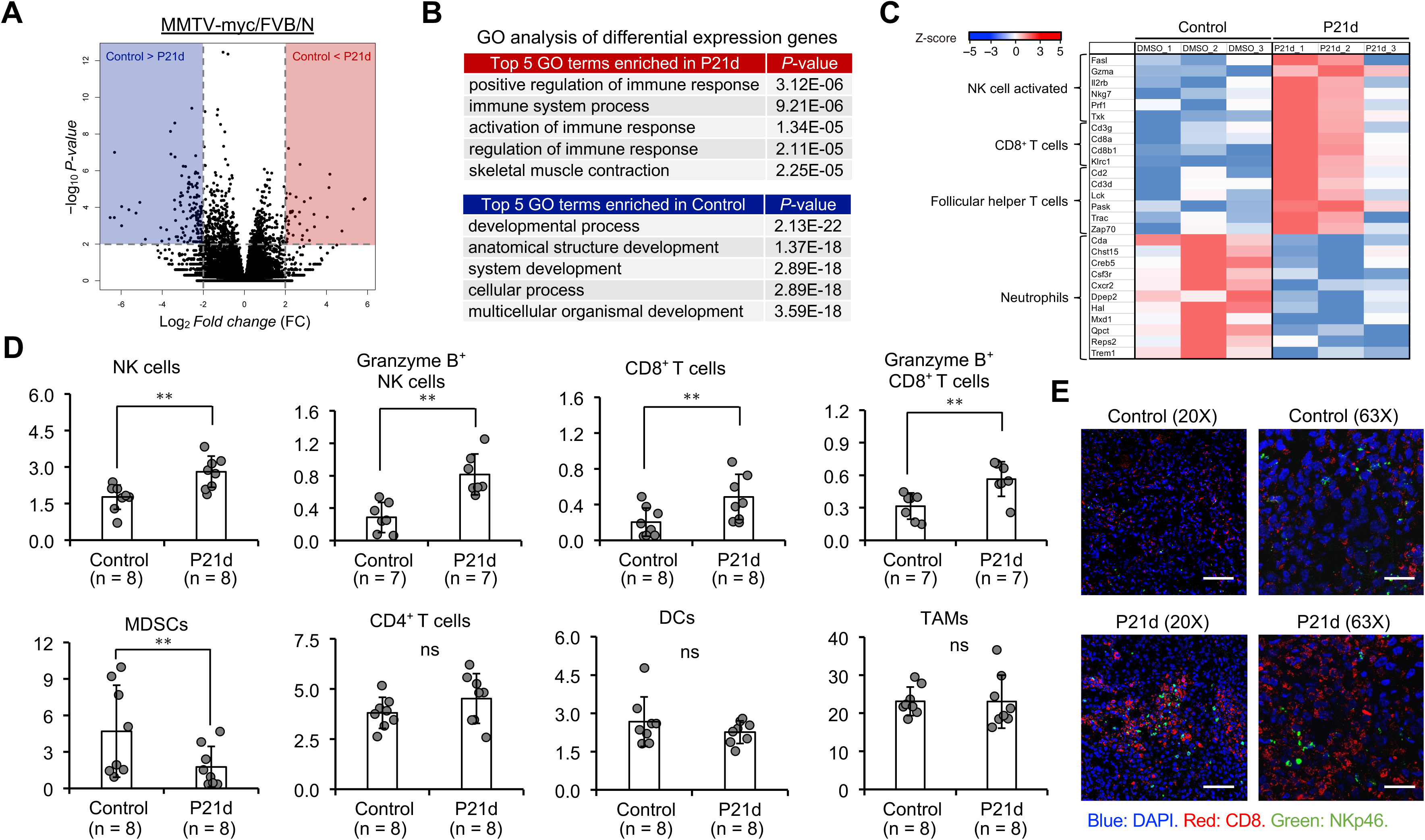
P21d induces a cytotoxic immune response in TNBC. (A) Volcano plot of genes differentially expressed in MMTV-myc tumors implanted in FVB/N mice treated with vehicle or P21d (30mg/kg/d) for 7 days. (B) Gene ontology (GO) analysis of upregulated and downregulated differentially expressed genes in MMTV-myc tumors treated with P21d relative to vehicle control-treated tumors. (C) Heatmap showing normalized expression levels in MMTV-myc tumors treated with vehicle or P21d for immune cell marker genes selected by CIBERSORTx. (D) Flow cytometry analysis of tumor infiltrating immune cells recovered from MMTV-myc primary tumors in FVB/N mice treated with vehicle or P21d (30mg/kg/d) for 7 days (n=7-8 mice /treatment group). Experiment was done three times. (E) Representative immunofluorescence images of MMTV-myc tumors treated with vehicle or P21d (30mg/kg/d) for 7 days and stained with antibody against CD8 or NKp46. Scale bars represent 200 μm (20X images) and 50 μm (63X images). Data are the mean ± SD. \*\**p*=0.01, \*\*\**p*=0.0001 by Student’s *t*-test.

We determined the immune cell types associated with the DEGs by examining immune cell marker genes. We defined immune cell marker genes based on the reference gene signature matrix used in CIBERSORT (See Methods for details) and performed enrichment tests between each immune cell marker and differentially expressed genes. We observed that activated NK and CD8^+^ T cell-related genes were highly expressed in P21d-treated tumors (**Fig.2C**). Consistent with this, we observed significant increases in tumor-infiltrating NK and CD8^+^ T cells in P21d-treated tumors, as analyzed by flow cytometry (**Fig. 2D**; **gating strategy in Supplemental Fig S3**). The increase in tumor-infiltrating cytotoxic immune cells induced by P21d treatment was not a consequence of generalized immune stimulation as P21d treatment of non-tumor bearing mice did not significantly alter the immune cell populations in the spleen or peripheral blood (**Supplemental Fig S4**).

These tumor-infiltrating NK and CD8^+^ T cells were cytotoxic, as evidenced by granzyme B positivity. These CD8^+^ T and NK cells were found to infiltrate the tumor core, as assessed by immunofluorescence (**Fig. 2E**). In contrast, there was a decrease in immunosuppressive myeloid-0000derived suppressor cells (MDSCs), with no significant change in CD4^+^ T cell, dendritic cell or macrophage populations (**Fig. 2 D**).

These results indicated that PTK6 inhibitor treatment increased the recruitment of tumor-infiltrating cytotoxic immune cells which could be responsible for TNBC tumor growth inhibition in immunocompetent models.

### PTK6 shRNA inhibits tumor growth and increases tumor infiltrating cytotoxic immune cells

Given that systemic administration of P21d makes it challenging to distinguish tumor-intrinsic vs. tumor microenvironmental vs. systemic effects, we utilized tumor cells expressing PTK6 shRNA to determine whether PTK6 inhibition only in the tumor cell compartment phenocopies the effects of P21d on tumor growth and TIL recruitment. The results of these studies would help pinpoint the mechanisms responsible for the observed immune modulation.

MMTV-myc cells expressing vector control or PTK6 shRNA were implanted into FVB/N mice and tumor growth was monitored. Tumoral PTK6 downregulation strongly inhibited tumor growth (**Fig. 3A-C**). TILs recovered from control or PTK6 shRNA-expressing tumors showed increased Granzyme B^+^ NK and CD8^+^ T cells in PTK6-downregulated tumors (**Fig. 3D**, **gating strategy in Supplemental Fig S5**). In contrast, the percentage of MDSC’s was decreased in the PTK6-downregulated tumors compared to that in the control (**Fig. 3D**). Similar effects of PTK6 downregulation were also observed in triple negative 4T-1 tumors; there were increased numbers of tumor-infiltrating CD8^+^ T cells and decreased numbers of MDSCs. These results phenocopy the effects observed with P21d treatment, suggesting that the inhibition of PTK6 in tumor cells is critical for changes in TIL populations. Of note, in contrast to MMTV-myc tumors, there was no increase in 4T-1 tumor infiltrating NK cells (**Supplemental Fig S6**).

**Figure 3.**
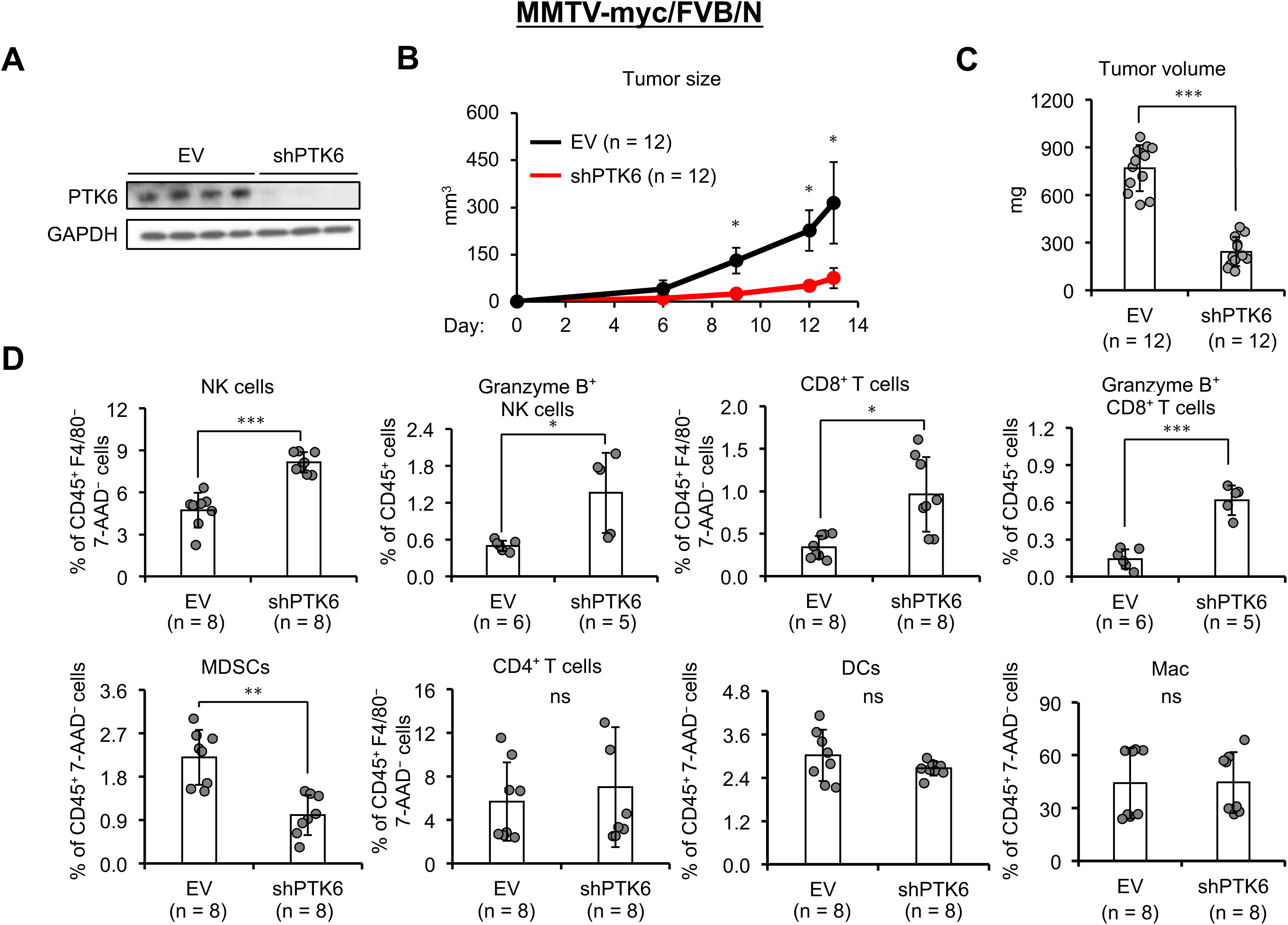
PTK6 shRNA inhibits TNBC growth and increases cytotoxic TILs. (A) PTK6 expression was assessed in primary tumors recovered from FVB/N mice 14 days after injection with MMTV-myc cells expressing either empty vector (EV) or PTK6 shRNA in the 4^th^ mammary fat pad. (n=3-4 mice/group). (B,C) FVB/N mice injected with MMTV-myc cells expressing EV or PTK6 shRNA were monitored for tumor growth (n=12 mice/group). At day 14, tumors from each group were recovered and weighed. (D) Flow cytometry analysis of tumor-infiltrating immune cells recovered from primary tumors expressing vector control or PTK6 shRNA (n=5-8/treatment group). Recovered TILs were stained with indicated antibodies and analyzed. Experiment was done three times. Data are the mean ± SD. \**p*=0.05, \*\**p*=0.01, \*\*\**p*=0.0001 by Student’s *t*-test.

### PTK6 inhibitor-induced increases in tumor infiltrating cytotoxic immune cells are important for tumor growth inhibition

CD8^+^ T cells and NK cells are critical for mobilizing adaptive and innate immunity, respectively, for the immunological control of cancers [32, 33]. We wondered whether the increased numbers of tumor-infiltrating CD8^+^ T cells and NK cells induced by P21d treatment mechanistically contribute to tumor growth inhibition observed with P21d treatment. If they contribute to a critical mechanism, we reasoned that the depletion of CD8^+^ T cells and/or NK cells might compromise P21d-dependent tumor growth inhibition. We confirmed by flow cytometry that CD8 antibody (Clone 53-6.7) and NK1.1 antibody (Clone PK136) depleted the intended immune populations (**Supplemental Fig S7A**). MMTV-myc tumor-bearing FVB/N mice were treated with either CD8 or NK1.1 antibodies prior to the initiation of P21d treatment. Antibody treatment was continued concurrently with P21d treatment as indicated (**Fig. 4**). Depletion of either CD8^+^ T cells or NK cells almost completely reversed the tumor growth inhibitory effect of P21d treatment (**Fig. 4**). As expected, depletion antibody treatment eliminated the targeted cells from the TIL populations (**Supplemental Fig S7B**). These results provide further evidence that immune microenvironmental regulation is a critical mechanism responsible for P21d-dependent inhibition of tumor growth. These results also indicate that both CD8^+^ T cell and NK cell activities are required for tumor inhibition, as depletion of either nearly completely abrogates P21d’s effects.

**Figure 4.**
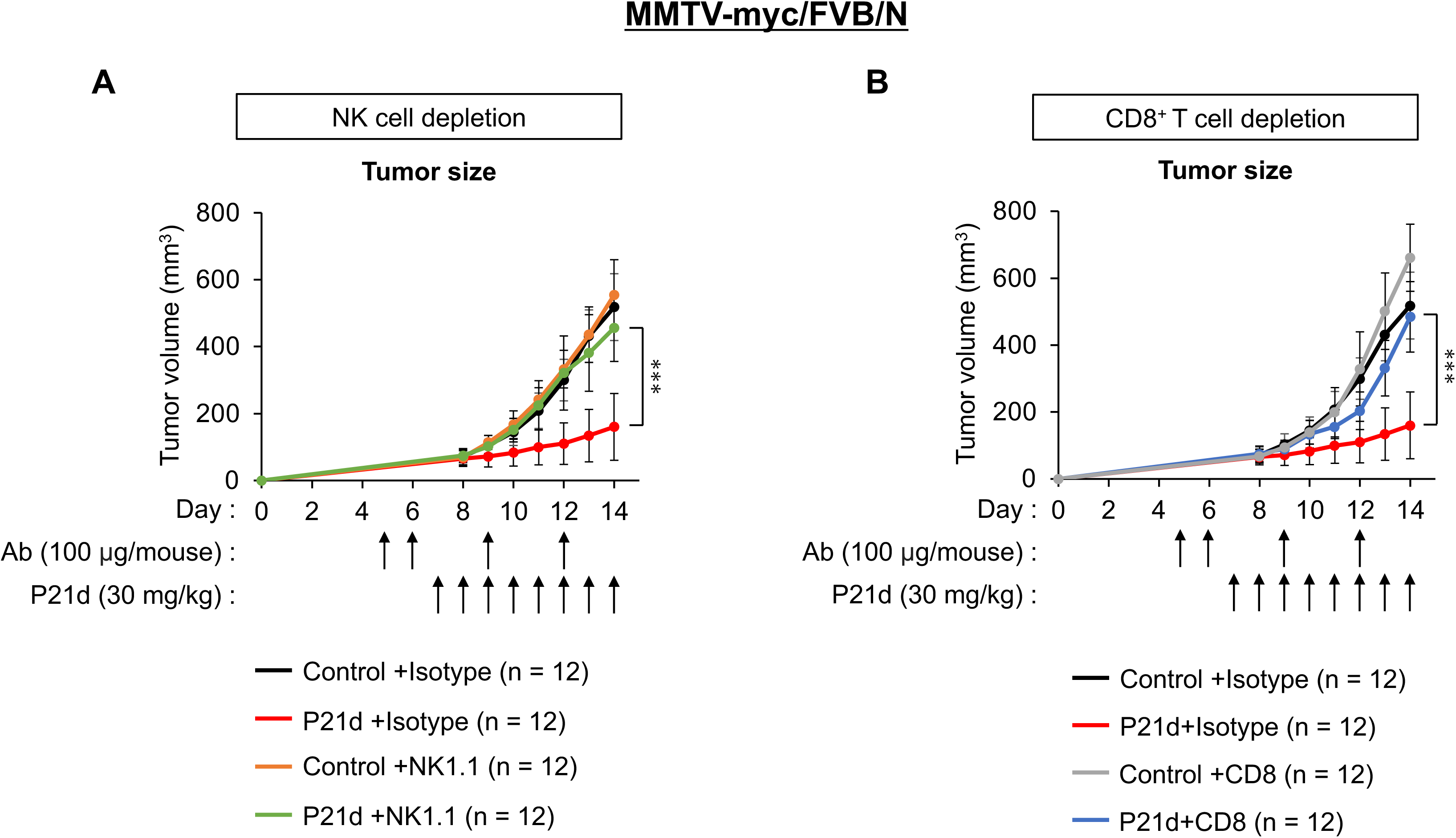
CD8^+^ T cell or NK cell depletion impairs P21d’s tumor inhibitory effects. (A) MMTV-myc tumor bearing FVN/N mice were pre-/co-treated with isotype or anti-NK1.1 antibody (100 μg/mouse) and vehicle or P21d (30mg/kg) (n=12 mice/treatment group). Primary tumor growth was measured on the indicated days. (B) MMTV-myc tumor bearing FVN/N mice were co-treated with isotype or anti-CD8 antibody (100 μg/mouse) and vehicle or P21d (30mg/kg) (n=12 mice/treatment group). Primary tumor growth was measured on the indicated days. Data are the mean ± SD. \*\*\**p*=0.0001 by Student’s *t*-test.

### SNAIL down-regulation inhibits tumor growth and phenocopies immunomodulatory effects of PTK6 inhibition

We have previously reported SNAIL as a downstream target of PTK6 [16, 19]. Mechanistically, PTK6 inhibition promotes SNAIL degradation by increasing its interaction with, and subsequent ubiquitination by MARCH2 E3 ligase [19]. SNAIL also promotes EMT-dependent tumor immunosuppression via multiple mechanisms. For example, SNAIL suppresses tumor immunity by regulating the expression of cytokines and chemokines, such as TSP1, and by regulating recruitment and function of MDSC’s and tumor associated macrophages (TAM) [10, 21]. To determine if SNAIL regulation mechanistically contributes to P21d’s immunomodulatory effects, we assessed whether downregulating tumoral SNAIL phenocopies the effects of P21d treatment. MMTV-myc cells expressing SNAIL shRNA were implanted into FVB/N mice and tumor growth was monitored (**Fig. 5A-C**). Downregulation of SNAIL inhibits tumor growth. In addition, flow cytometric analysis revealed an increase in tumor-infiltrating CD8^+^ T and NK cells in SNAIL shRNA-expressing tumors, with a decrease in infiltrating MDSCs (**Fig. 5D**, **gating strategy in Supplemental Fig. S8**). These results phenocopy the effects of P21d and support SNAIL regulation as a mediator of P21d’s immunoregulatory functions, as well as the importance of EMT regulation in tumor immune microenvironment composition.

**Figure 5.**
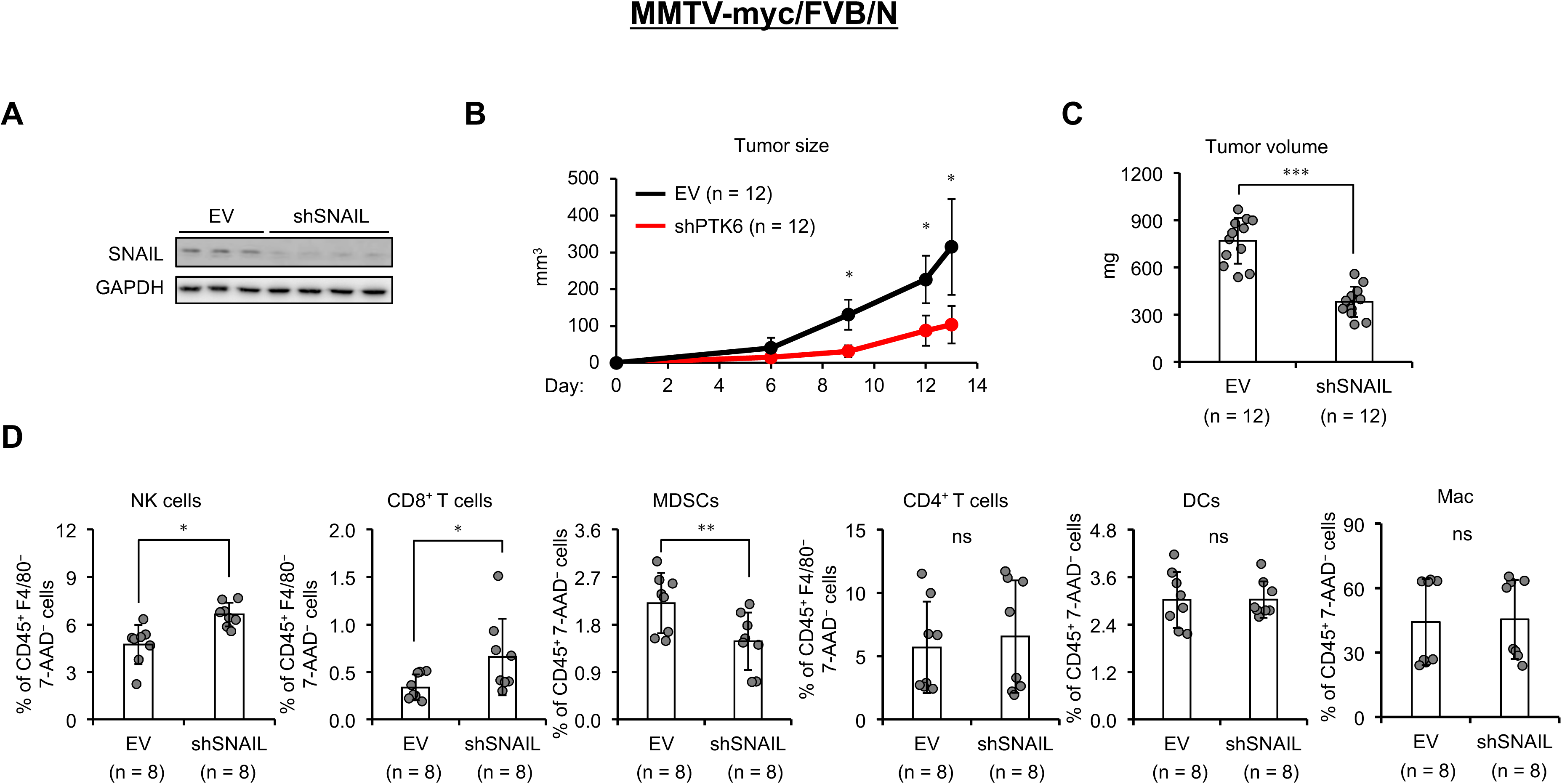
SNAIL downregulation inhibits TNBC tumor growth and increases cytotoxic TILs. (A) SNAIL expression in primary tumors generated by injecting FVB/N mice with MMTV-myc cells expressing empty vector (EV) or Snail shRNA. (B,C) FVB/N mice injected with MMTV-myc cells expressing EV or Snail shRNA were monitored for tumor growth (n=12 mice/group). At day 14, tumors from each group were recovered and weighed. (D) Flow cytometry analysis of tumor-infiltrating immune cells recovered from primary tumors expressing vector control or Snail shRNA (n=8/treatment group). Recovered TILs were stained with indicated antibodies and analyzed. Experiment was done three times. Data are the mean ± SD. \**p*=0.05, \*\**p*=0.01 by Student’s *t*-test.

### PTK6 inhibition increases TIL recruitment via CXCL10/CXCR3

We wondered if the increase in cytotoxic TILs observed in P21d treated mice could be due to the increased recruitment of these cells to the tumor site. We tested whether P21d treatment of tumor cells produces factors that can act as chemoattractants for immune cell migration and eventual tumoral infiltration. Conditioned media (culture supernatants) from MMTV-myc cells treated *in vitro* with either vehicle control or P21d were applied to the bottom chamber of the Transwell assay units. CD8^+^ T cells or NK cells isolated from mouse spleen were applied to the top chamber and the number of migrated immune cells was counted. Conditioned media from P21d-treated cells increased the migration of both CD8^+^ T and NK cells several-fold compared to that of vehicle-treated cells (**Fig. 6A**). These results suggest that P21d treatment of tumor cells leads to the production of secreted factors that induce the migration of immune cells, which could be responsible, at least in part, for increased tumoral infiltration.

**Figure 6.**
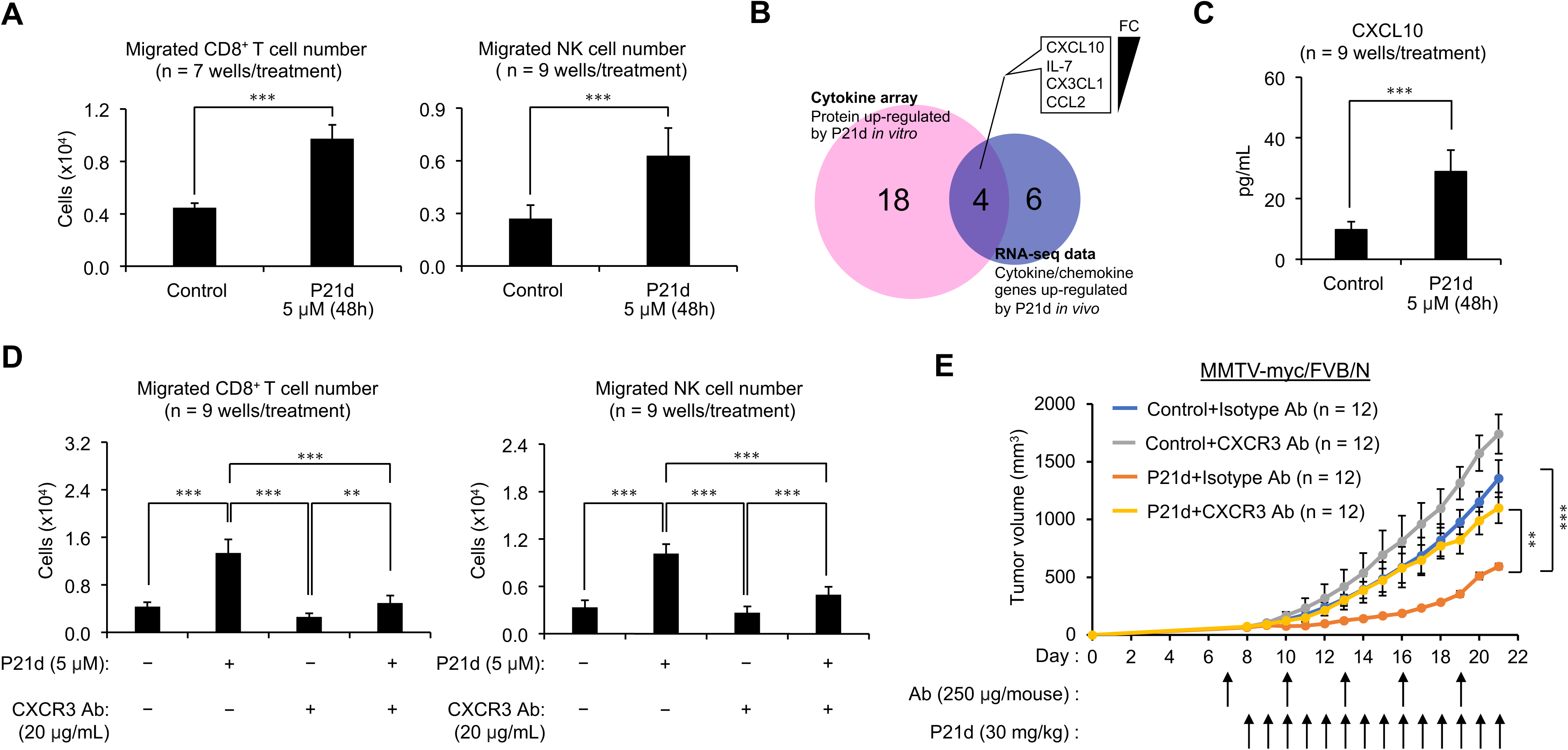
P21d induces tumoral secretion of CXCL10 to recruit cytotoxic TILs and inhibit TNBC tumor growth. (A) Migration assays were performed using NK cells and CD8^+^ T cells isolated from non-tumor bearing mouse spleens and supernatants (conditioned media) of MTV-myc cells treated *in vitro* for 48 hours with vehicle or P21d (5 μM) (n=7-9 wells/treatment condition). (B) A Venn diagram showing the overlap of molecules upregulated in both RNA sequencing analysis of MMTV-myc tumors treated *in vivo* with P21d (30mg/kg/mice for 7 days) and cytokine arrays performed using MMTV-myc cells treated *in vitro* with P21d (5μM) for 48 hours. (C) ELISA assay was performed to assess levels of CXCL10 in culture supernatants of MMTV-myc cells treated *in vitro* with vehicle or P21d (5μM) for 48 days (n=9 wells/treatment condition). (D) Transwell migration assays were performed using NK cells or CD8^+^ T cells isolated from non-tumor bearing mouse spleen. Isolated NK cells or CD8^+^ T cells were pre-treated with anti-CXCR3 antibody (20 μg/mL) or isotype control IgG for 30 minutes and subsequently applied to the top chamber in the continued presence of anti-CXCR3 or isotype control antibody. Conditioned medium from MMTV-myc cells treated *in vitro* for 48 hours with vehicle control or P21d (5μM) was applied to the bottom chamber. Migration was assessed after 4 hours (n=9 wells/treatment condition). (E) MMTV-myc tumor bearing FVB/N mice were pre-/co-treated with isotype IgG or anti-CXCR3 antibody (250 μg/mouse) and vehicle or P21d (30mg/kg/d) (n=12 mice/treatment group). Primary tumor growth was measured on the indicated days. Data are the mean ± SD. \**p*=0.05, \*\**p*=0.01, \*\*\**p*=0.0001 by Student’s *t*-test.

To identify mediators of this chemotactic response, we analyzed data from RNA-sequencing of MMTV-myc tumors treated *in vivo* with P21d. We also performed cytokine arrays using the supernatants of cells treated *in vitro* with P21d (**Supplemental Fig S9A**). We focused on candidates that were upregulated by P21d treatment in both studies relative to control. Among them, CXCL10 was associated with the greatest fold-change relative to control (**Fig. 6B**, **Supplemental Fig. S9A**). CXCL10 is produced by both immune cells and cancer cells. CXCR3, the receptor for CXCL10, is highly expressed on T and NK cells, and CXCL10 is known to recruit T cells and NK cells [34]. We focused on CXCL10 and performed ELISA assays to confirm the increase in secreted CXCL10 in the supernatants of P21d-treated cultures (**Fig. 6C**).

Next, we sought to determine whether the increased level of CXCL10 induced by P21d treatment was functionally responsible for the enhanced chemotactic migration of immune cells. We assessed the effect of a blocking antibody against CXCR3, the receptor for CXCL10, on migration induced by conditioned media from P21d-treated cells *in vitro*, as well as tumor infiltration by NK or CD8^+^ T cells *in vivo*. Co-incubation with the anti-CXCR3 antibody inhibited the enhanced chemotactic migration of NK and CD8^+^ T cells in Transwell assays previously observed with P21d conditioned media (**Fig. 6D**). Similarly, co-treatment of MMTV-myc tumor-bearing mice with anti-CXCR3 antibody and P21d reduced tumoral infiltration by NK and CD8^+^ T cells (**Supplemental Fig. S9B**). In addition, anti-CXCR3 antibody/P21d co-treatment almost completely reversed the tumor growth inhibitory effects observed with P21d treatment (**Fig. 6E**). These data again strongly support the importance of TIL recruitment for tumor growth inhibition and that this recruitment occurs via CXCL10/CXCR3 signaling.

## DISCUSSION

EMT contributes to tumor immune evasion and immunosuppression through multiple mechanisms, enabling cancer cells to bypass the cytotoxic activity of immune cells that helps prevent dissemination and metastases [22, 35, 36]. Therefore, targeting EMT drivers is an attractive strategy to favorably remodel the tumor immune microenvironment, boost anti-tumor immune responses and potentially improve the response to immunotherapies. The present study highlights the immunological consequences of inhibiting PTK6, a targetable driver of EMT, as well as its downstream effector SNAIL, which could be leveraged to promote a more robust immune response against TNBC.

Cancer cells that have undergone EMT are more able to evade immune recognition and killing because of the downregulation of antigen-presenting molecules such as MHC Class I and the increased expression of immunosuppressive molecules such as PD-L1, CD73, and CD276 (reviewed in [14, 37]). In addition, mesenchymal cells secrete factors such as TGFβ, CSF1, CCL18 and CXCL1/2, which recruit immunosuppressive regulatory T cells (Tregs), tumor-associated macrophages (TAMs) and myeloid-derived suppressor cells (MDSCs) that can dampen effective anti-tumor immune responses [11, 13, 35, 38–40]. Despite overwhelming evidence supporting the inhibitory effects of EMT on cytotoxic CD8^+^ T cell responses, the effects of EMT on NK cytotoxic responses have been somewhat more controversial and may depend on specific stages of tumor progression and metastases, as well as the balance in EMT-induced expression of molecules that interact with activating or inhibitory NK cell receptors. For example, EMT-associated downregulation of MHC Class I and E-cadherin, which function as inhibitory ligands of NK cells, renders tumor cells more susceptible to NK cell-mediated cytotoxicity [41]. However, the expression of other molecules, such as galectin 3 and ligands for inhibitory NK cell receptors such as HLA-G and HLA-E, is also induced during EMT leading to the suppression of NK cell function [42].

Transcriptional drivers of EMT, such as SNAIL, SLUG, TWIST, and Zeb1, have been directly linked to the production and secretion of chemokines such as CCL2, CXCL1/2, and CSF1 that recruit immunosuppressive Tregs and MDSCs [20, 43–45]. However, these transcription factors have been challenging to target directly and there are no clinically translatable candidates. Therefore, targeting the regulators of these transcription factors may offer an alternative therapeutic approach. We previously reported that PTK6 is expressed at high levels in approximately 40% of TNBC cases, and its increased expression is sufficient to drive EMT by promoting the stabilization of SNAIL [16, 19]. PTK6 expression in patient triple negative tumors is also correlated with SNAIL levels, and PTK6 inhibition downregulates SNAIL levels by promoting its ubiquitination by MARCH2 E3 ligase and proteasome-dependent degradation [19]. Thus, targeting PTK6, downregulating SNAIL, and blocking or reversing EMT could enhance immune responses against a subset of TNBC.

Inhibition of PTK6 kinase activity with P21d treatment increased the recruitment of both cytotoxic CD8^+^ T and NK cells. There was an increase in Granzyme B positive cells, indicating the cytotoxic activity of these TILs. Simultaneously, tumor-infiltrating immunosuppressive MDSCs were decreased, providing additional support for favorable remodeling of the tumor immune microenvironment. The effects of P21d treatment on TILs were phenocopied by downregulating PTK6 expression in tumor cells using PTK6 shRNA, suggesting that inhibition of PTK6 within the tumor cell compartment is sufficient to increase cytotoxic TILs. Although we did not observe any dramatic changes in immune populations in the peripheral blood or spleens of P21d-treated mice, we cannot completely rule out a complementary contribution of the systemic immune effects of P21d treatment, especially in light of recent reports of PTK6’s functions as an immune modulator [46–48]. A germline inactivating PTK6 mutation is associated with rare cases of familial lupus, an autoimmune condition [46, 47]. This mutation is located in the kinase domain and leads to decreased PTK6 activity. Therefore, decreased PTK6 activity in systemic immune cells could contribute to a heightened response. In addition, PTK6 plays a role in intestinal type 2 innate immune responses by regulating IL-25 and IRAG2[48].

Interestingly, PTK6 was recently proposed as a biomarker for prognosis and response to immunotherapy for renal cell carcinoma [28]. Similar to breast cancer, higher levels of PTK6 expression were linked to a worse prognosis. PTK6 expression was also correlated with distinct immune signatures. In a pan-cancer analysis, Xiong et al. reported a positive correlation between PTK6 expression and immunosuppressive Treg cells, as well as a negative correlation between PTK6 and CD4+ and CD8^+^ T cells [49]. This study also established positive and negative correlations between PTK6 and various immunomodulatory chemokines.

We identified CXCL10 induction as a mechanism for P21d-mediated recruitment of CD8^+^ T cells and NK cells that are responsible for tumor growth inhibition. CXCL10 is secreted by a variety of cell types, including leukocytes, epithelial cells, fibroblasts and endothelial cells and contributes to inflammatory processes in infectious and autoimmune disease, as well as cancer [34, 50]. CXCL10 has been reported to play a pro-or anti-tumorigenic role, depending on the tumor type and expression patterns and levels of CXCR3, the receptor for CXCL10 [34]. CXCL10 is an established “homing factor” for NK cells, activated T cells, and antigen-presenting dendritic cells that help to enhance innate and adaptive immunity to contain tumor growth. CXCL10 also has an anti-angiogenic effect, inhibiting vascular endothelial growth factor levels and blocking the formation of microvessels to inhibit the growth of cervical and ER^+^ breast cancers [51, 52]. In contrast, autocrine activation of CXCR3 expressed at high levels on tumor cells has been reported to promote tumor progression and metastasis [53, 54]. In our studies co-treatment with a blocking CXCR3 antibody restored tumor growth of P21d-treated TNBC, which does not support autocrine pro-tumorigenic CXCL10:CXL10 signaling as a dominant mechanism in this model.

These effects of PTK6 inhibition on the immune microenvironment of TNBC also support targeting EMT drivers, such as PTK6, as an immunotherapy-sensitizing strategy. While immunotherapy is the approved standard of care for stage 2 and 3 triple negative breast cancer, responses vary and there is a need for strategies to improve response. Ongoing studies to assess the effects of PTK6 inhibition on checkpoint and other immunoregulatory molecules should provide insights into optimal combination therapies.

In conclusion, PTK6 kinase inhibition upregulates the tumoral expression of the CXCR3 ligand CXCL10 and enhances CD8^+^ T cells and NK cells tumor infiltration via CXCR3 engagement, enhancing anti-tumor immunity to suppress tumor growth. These findings provide new insights into the mechanisms linking EMT to tumor immune responses. Thus, PTK6 kinase inhibition could be an effective therapeutic strategy in combination with immunotherapies for a subset of TNBC.

## Competing Interests

The authors declare no potential conflicts of interest.

## Acknowledgments

The authors acknowledge the Tisch Cancer Institute Biorepository and Histopathology Cores at the Icahn School of Medicine at Mount Sinai. This work was supported by a Research Scholar Grant (18-044-01-LIB) and an Accelerator Award (EDS-24-1331013-01-EDS) from the American Cancer Society to H.Y.I. H.Y.I. was also supported by a grant from the Breast Cancer Research Foundation (22-130). K.I. was supported by a Post-doctoral Fellowship from Susan G. Komen for the Cure.

## Author Contributions

Conceptualization: I.H., C.M., K.I. and H.Y.I.; Methodology: I.H., C.M., K.I., E.L., J.Z., and H.Y.I.; Formal Analysis: I.H., C.M., K.I., E.L., J.Z., and H.Y.I ; Investigation: I.H., C.M., K.I., E.L., J.Z., and H.Y.I.; Writing: I.H., C.M., E.L., J.Z., and H.Y.I.; Supervision and Funding Acquisition: K.I. and H.Y.I.

## Data Availability Statement

The datasets generated during and/or analyzed during the current study are available from the corresponding author upon reasonable request.

## Notes

### Competing Interest Statement

The authors have declared no competing interest.

