## Supplemental Data for "Inhibition of EMT driver PTK6 enhances anti-tumor immune responses against triple-negative breast cancer"

Supplemental Fig S1

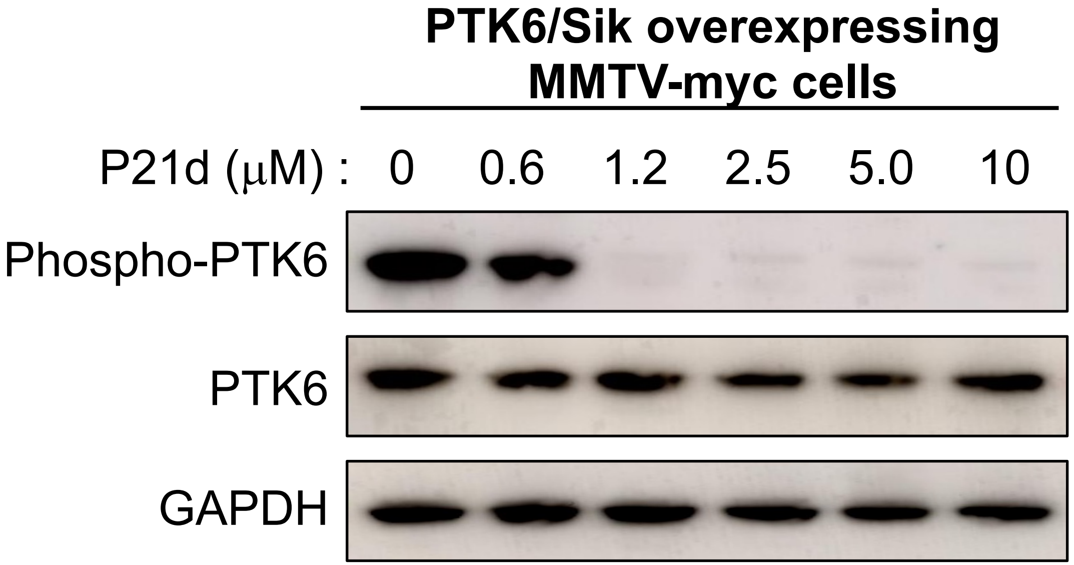

**Supplemental Figure Legends:**

**Supplemental Figure 1. P21d inhibits PTK6/Sik activity, as assessed by PTK6 autophosphorylation**

MMTV-myc cells stably overexpressing wild-type Sik (mouse PTK6) were treated with P21d at the indicated concentration for 3 hours, lysed and analyzed by Western blot using antibodies against total or autophosphorylated (Y342) PTK6.

### Supplemental Fig S2

#### Levels of P21d in MMTV-myc tumors

| | p21d: Summary of Tumor Levels | |
|---|---|---|
| | Sample ID in Raw Data | Calc Conc (ng/g) |
| NSG background | Am1_Tumor DMSO | BQL |
| | Am2_Tumor P21d | 4290 |
| | Am3_Tumor P21d | 4180 |
| | Am4_Tumor P21d | 5070 |
| FVB/N background | Am5_Tumor DMSO | BQL |
| | Am6_Tumor P21d | 100 |
| | Am7_Tumor P21d | 460 |
| | Am8_Tumor P21d | 544 |

BQL = Below Quantitation Limit of 0.400 ng/mL (or 20 ng/g after 10x homogenization dilution and 5x assay dilution)

**Supplemental Figure 2. Intratumoral concentration of P21d in MMTV-myc tumors implanted in FVB/N or NSG mice.**

Tumor bearing mice were treated with P21d (30mg/kg/d) for 3 days. Tumors were recovered and analyzed by mass spectrometry (Charles River Labs)

### Supplemental Fig S3

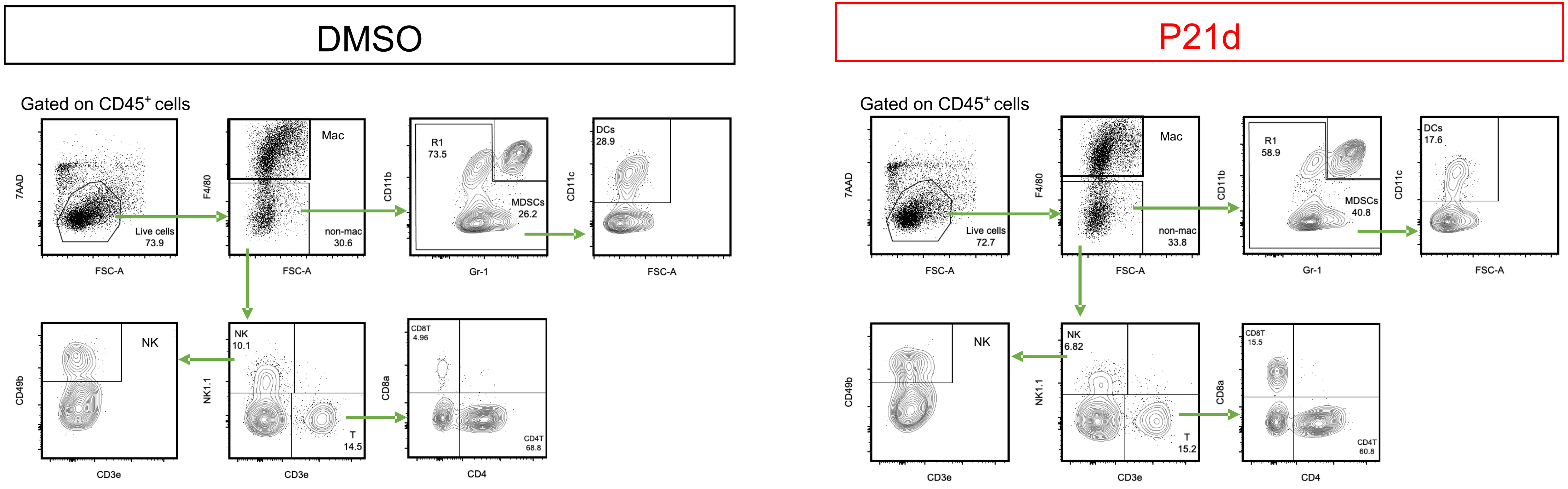

**Supplemental Figure 3. Flow cytometry gating strategy**

Gating strategy for flow cytometry analysis of TILS. In this analysis, $CD45^{+}$ 7-$AAD^{-}$ $F4/80^{+}$ cells are tumor-associated macrophages (TAMs), $CD45^{+}$ 7-$AAD^{-}$ $F4/80^{-}$ $CD11b^{+}$ Gr-$1^{+}$ cells are myeloid derived suppressor cells (MDSCs), $CD45^{+}$ 7-$AAD^{-}$ $F4/80^{-}$ $CD11c^{+}$ cells are dendritic cells (DCs), $CD45^{+}$ 7-$AAD^{-}$ $F4/80^{-}$ $CD3e^{-}$ $NK1.1^{+}$ $CD49b^{+}$ cells are NK cells, $CD45^{+}$ 7-$AAD^{-}$ $F4/80^{-}$ $NK1.1^{-}$ $CD3e^{+}$ $CD4^{-}$ $CD8a^{+}$ cells are $CD8^{+}$ T cells and $CD45^{+}$ 7-$AAD^{-}$ $F4/80^{-}$ $NK1.1^{-}$ $CD3e^{+}$ $CD4^{+}$ $CD8a^{-}$ cells are $CD4^{+}$ T cells.

#### Supplemental Fig S4

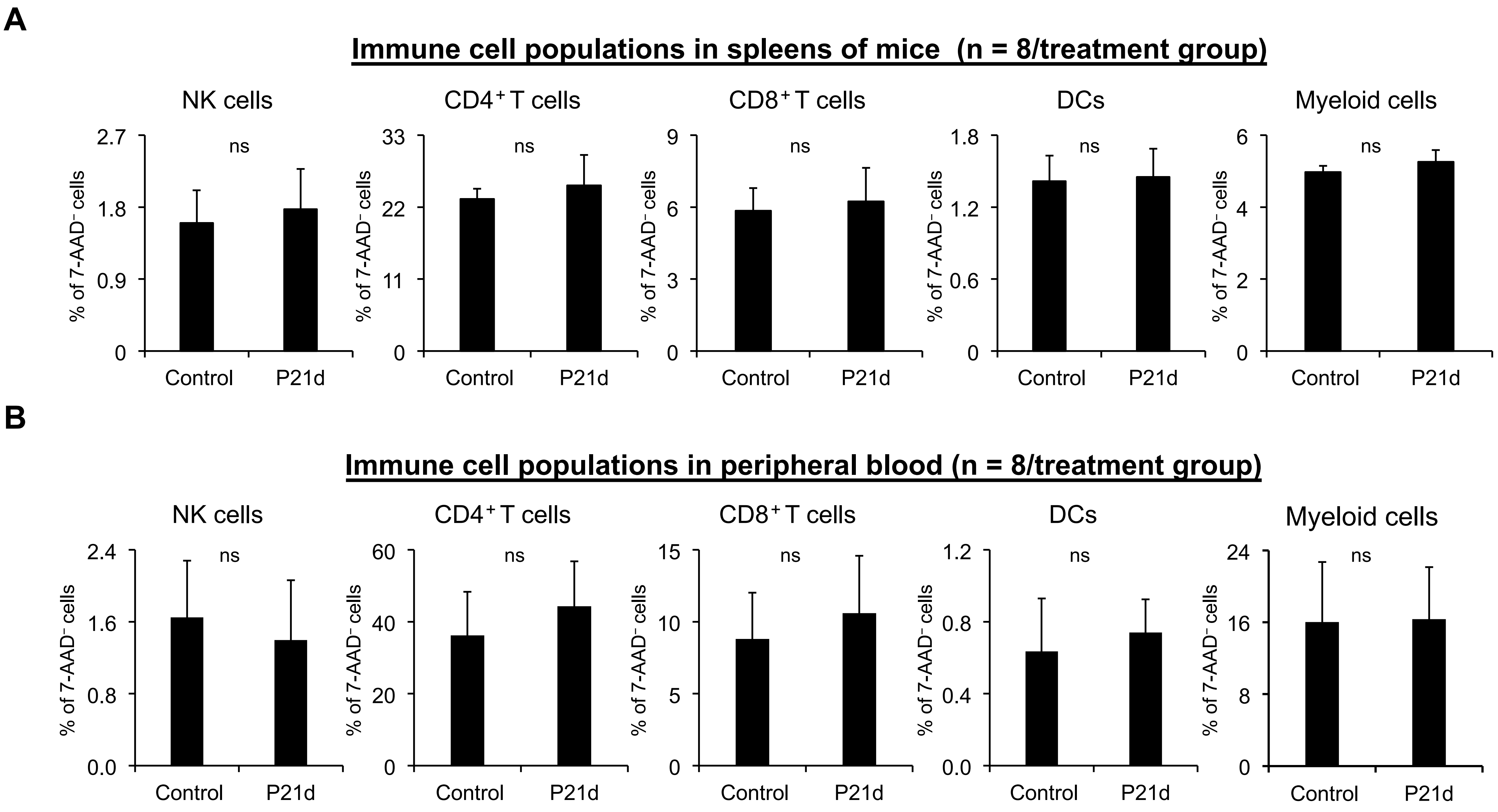

**Supplemental Figure 4. P21d does not affect immune cells in peripheral blood or spleen**

Immune cells ($CD8^+$ T cells, $CD4^+$ T cells, NK cells, Myeloid cells, DCs) in spleen and peripheral blood were isolated from non-tumor bearing mice treated with vehicle control or P21d (30mg/kg/d) for seven days (n=8 mice/treatment group).

### Supplemental Fig S5

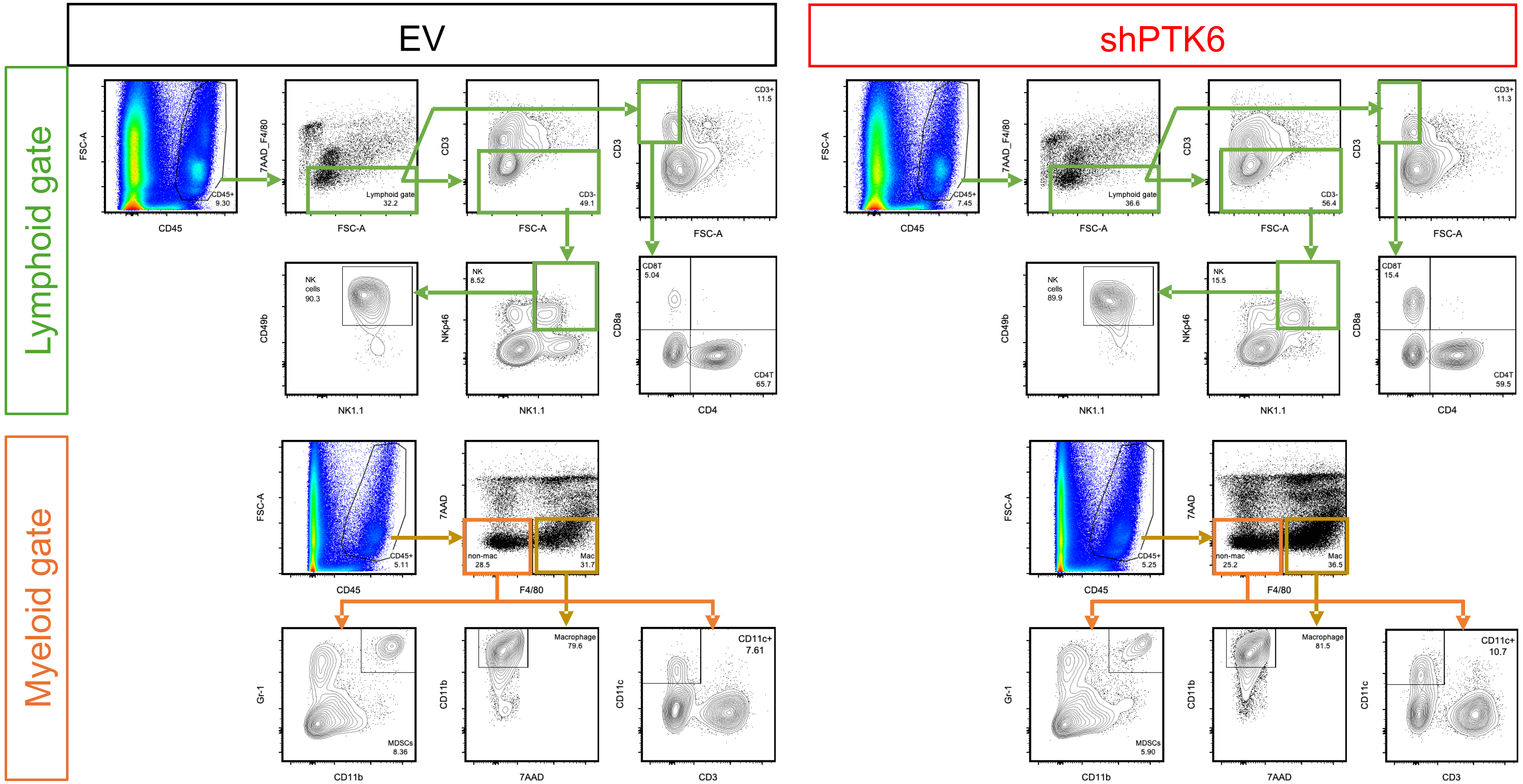

**Supplemental Figure 5. Flow cytometry gating strategy**

Gating strategy for flow cytometric analysis of TILs. $CD45^{+}$ $7\text{-}AAD^{-}$ $F4/80^{+}$ $CD11b^{+}$ cells are tumor-associated macrophages (TAMs), $CD45^{+}$ $7\text{-}AAD^{-}$ $F4/80^{-}$ $CD11b^{+}$ $Gr\text{-}1^{+}$ cells are myeloid derived suppressor cells (MDSCs), $CD45^{+}$ $7\text{-}AAD^{-}$ $F4/80^{-}$ $CD11c^{+}$ cells are dendritic cells (DCs), $CD45^{+}$ $7\text{-}AAD^{-}$ $F4/80^{-}$ $CD3e^{-}$ $NK1.1^{+}$ $CD49b^{+}$ $NKp46^{+}$ cells are NK cells, $CD45^{+}$ $7\text{-}AAD^{-}$ $F4/80^{-}$ $NK1.1^{-}$ $CD3e^{+}$ $CD4^{-}$ $CD8a^{+}$ cells are $CD8^{+}$ T cells and $CD45^{+}$ $7\text{-}AAD^{-}$ $F4/80^{-}$ $NK1.1^{-}$ $CD3e^{+}$ $CD4^{+}$ $CD8a^{-}$ cells are $CD4^{+}$ T cells.

### Supplemental Fig S6

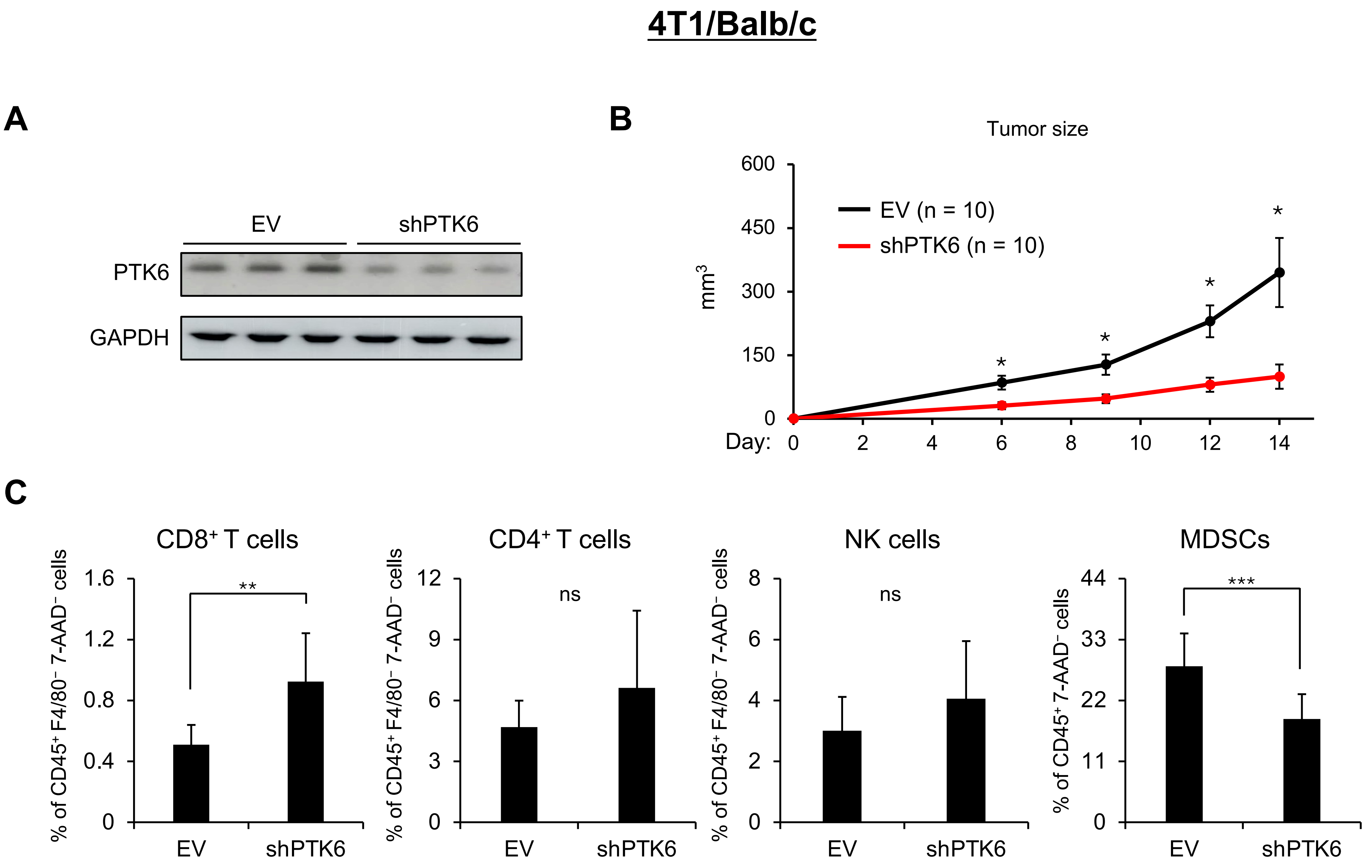

**Supplemental Figure 6. PTK6 shRNA inhibits 4T-1 tumor growth and increases TILs.**

(A) PTK6 expression in primary tumors generated by injecting Balb/C mice with 4T-1 cells expressing empty vector (EV) or PTK6 shRNA.

(B) Balb/C mice injected with 4T-1cells expressing EV or PTK6 shRNA were monitored for tumor growth (n=10 mice/group).

(C) Flow cytometry analysis of tumor-infiltrating immune cells recovered from primary tumors expressing vector control or Snail shRNA (n=10/treatment group). Recovered TILs were stained with indicated antibodies and analyzed.

Data are the mean ± SD. *p=0.05, **p=0.01, ***p=0.0001 by Student's t-test.

### Supplemental Fig S7

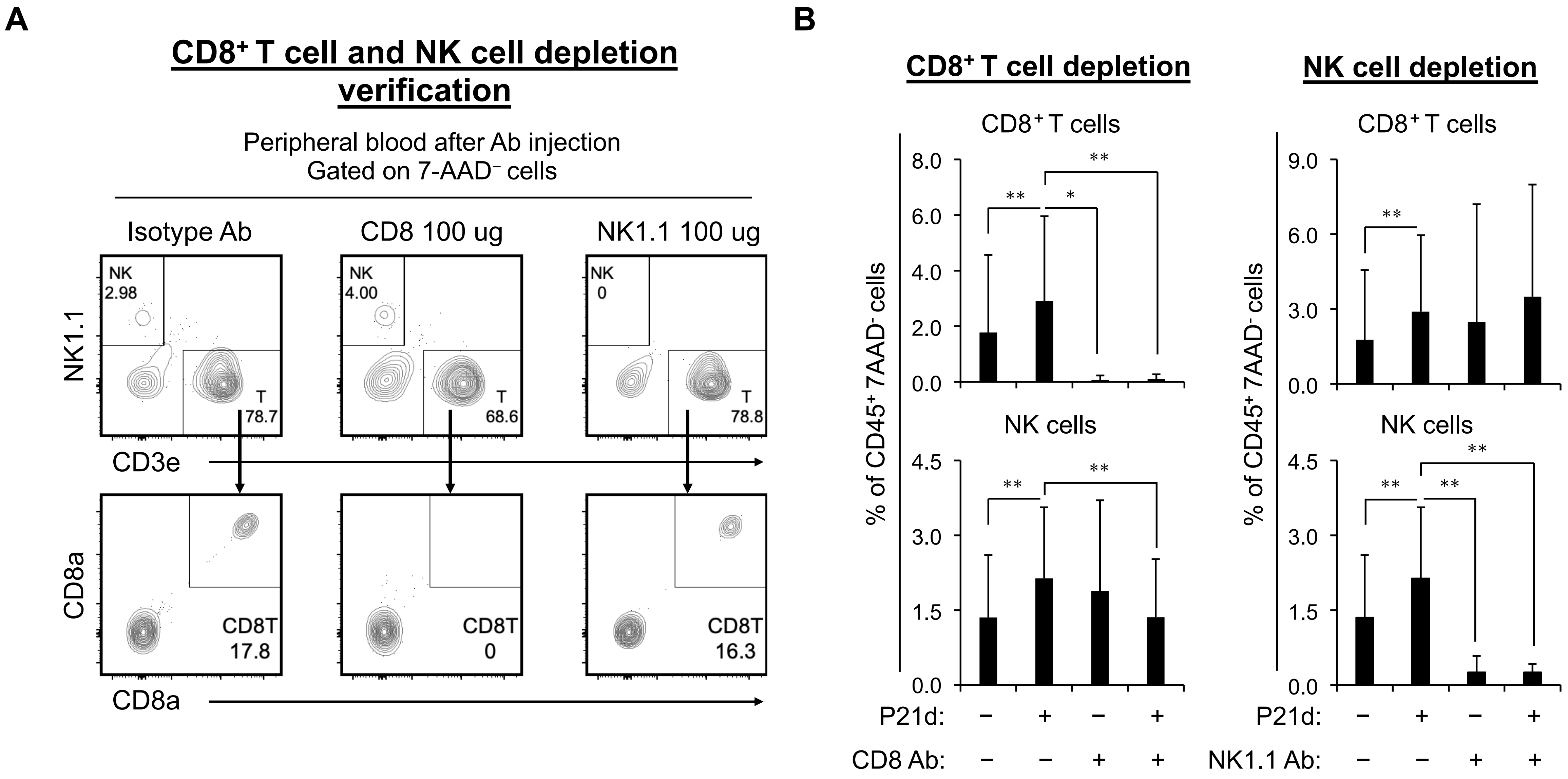

**Supplemental Figure 7. NK1.1 and CD8a antibodies effectively deplete NK and $CD8^+$ T cells**

(A) Representative FACS plots of NK cells and $CD8^+$ T cells in peripheral blood four days after single injection with anti-NK1.1 or CD8 antibody (n=3 mice/treatment group).

(B) Analysis of tumor-infiltrating NK cells and $CD8^+$ T cells in MMTV-myc tumor bearing mice treated with from NK1.1 or CD8 depleting antibody (100 μg/mouse) and vehicle control or P21d for seven days (n=6 mice/group).

Data are the mean ± SD. **p*=0.05, ***p*=0.01, ****p*=0.0001 by Student's *t*-test.

### Supplemental Fig S8

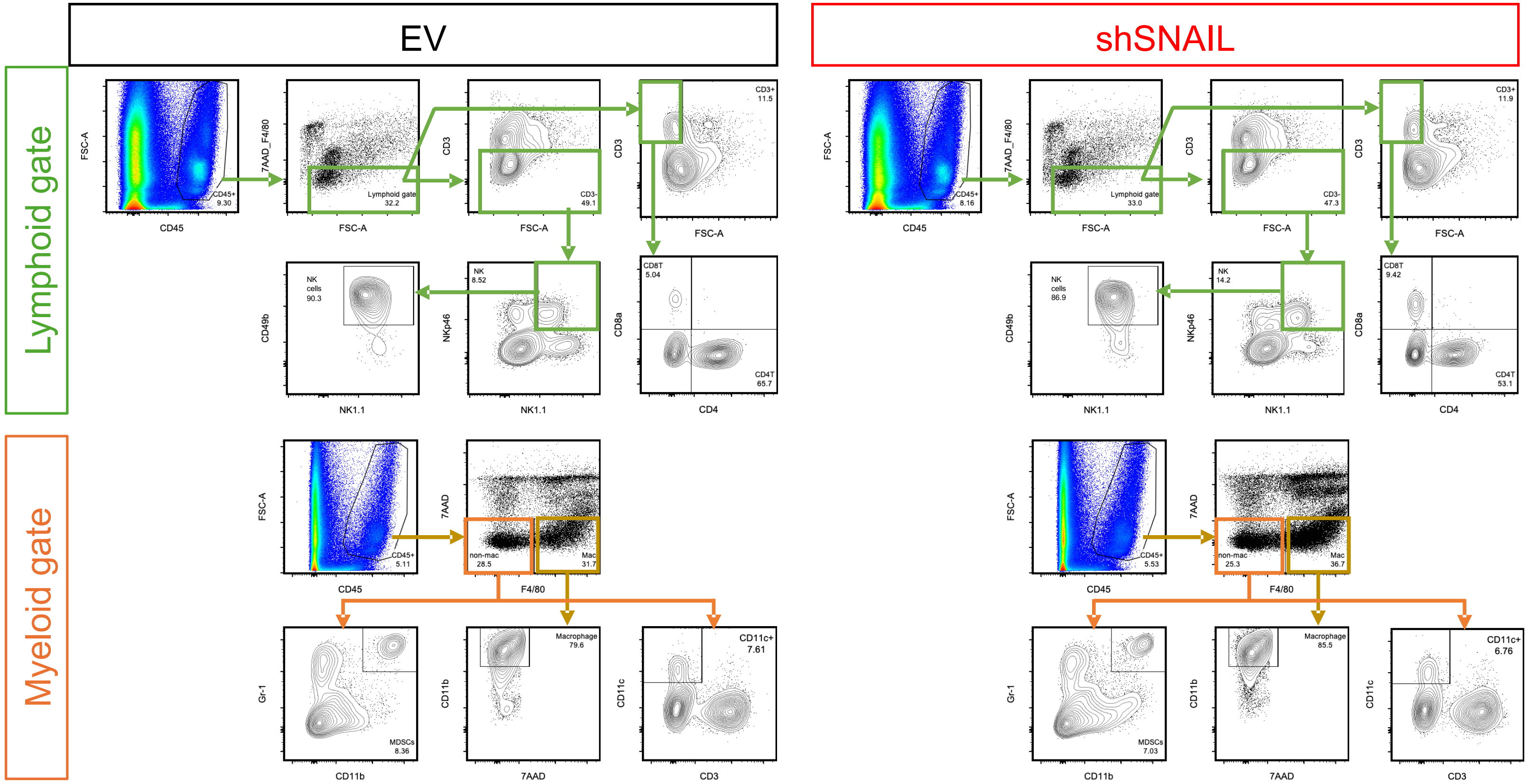

**Supplemental Figure 8. Flow cytometry gating strategy.**

Gating strategy: Representative FACS plots of in tumors (tumor infiltrating leukocyte analysis). In this analysis, $CD45^+$ 7-$AAD^-$ F4/$80^+$ $CD11b^+$ cells are tumor-associated macrophages (TAMs), $CD45^+$ 7-$AAD^-$ F4/$80^-$ $CD11b^+$ Gr-$1^+$ cells are myeloid derived suppressor cells (MDSCs), $CD45^+$ 7-$AAD^-$ F4/$80^-$ $CD11c^+$ cells are dendritic cells (DCs), $CD45^+$ 7-$AAD^-$ F4/$80^-$ $CD3e^-$ NK1.$1^+$ $CD49b^+$ $NKp46^+$ cells are NK cells, $CD45^+$ 7-$AAD^-$ F4/$80^-$ NK1.$1^-$ $CD3e^+$ $CD4^-$ $CD8a^+$ cells are $CD8^+$ T cells and $CD45^+$ 7-$AAD^-$ F4/$80^-$ NK1.$1^-$ $CD3e^+$ $CD4^+$ $CD8a^-$ cells are $CD4^+$ T cells.

#### Supplemental Fig S9

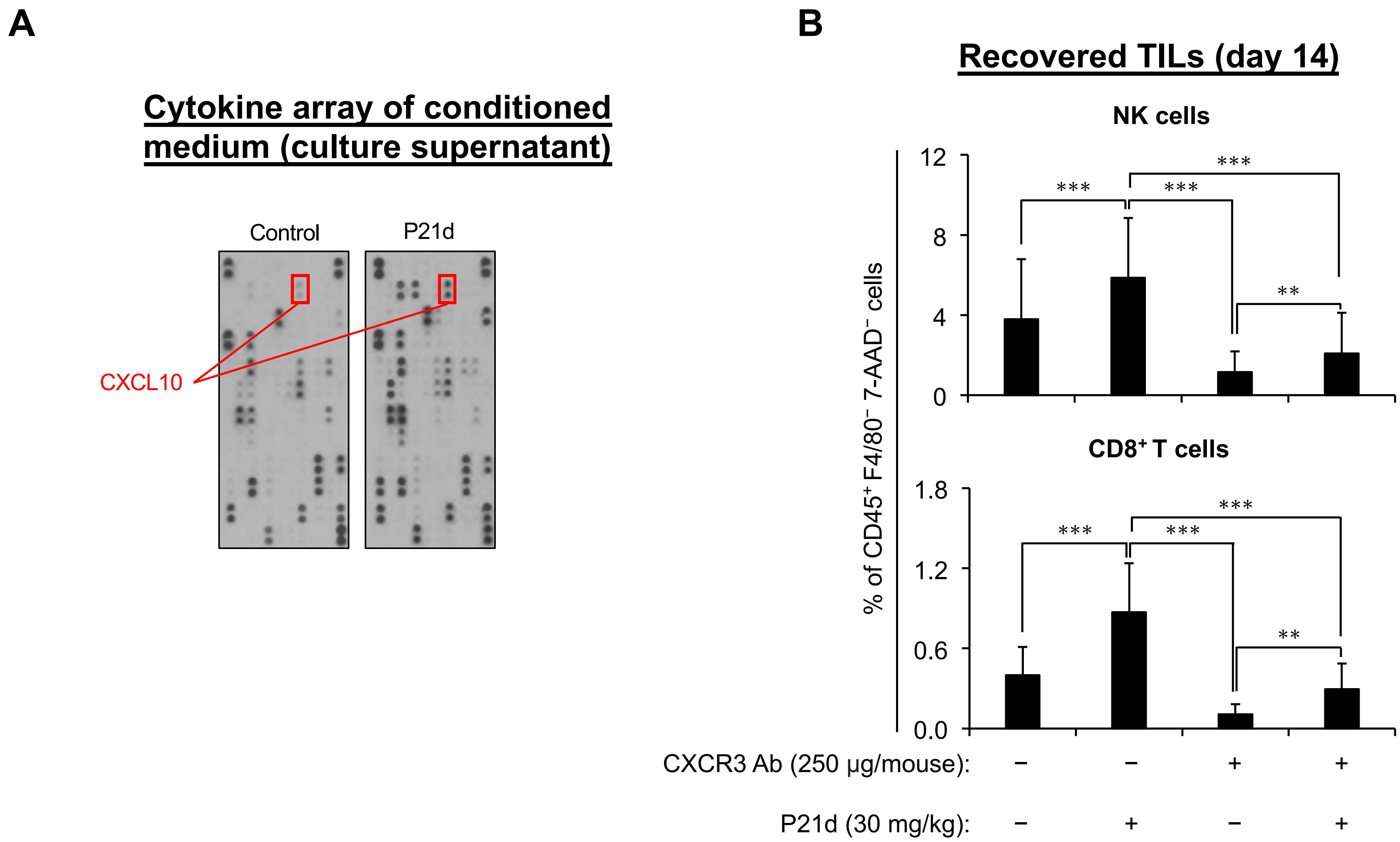

**Supplemental Figure 9. P21d induces CXCL10 production**

(A) Cytokine array analysis was performed using conditioned media from MMTV-myc cells treated with vehicle or P21d (5μM) for 48 hours.

(B) Flow cytometry analysis of tumor-infiltrating immune cells in MMTV-myc primary tumors recovered from FVB/N mice treated with anti-CXCR3 antibody and vehicle or P21d (30mg/kg) for 14 days (n=8 mice/treatment group).

Data are the mean ± SD. **p*=0.05, ***p*=0.01, ****p*=0.0001 by Student's *t*-test.
